# Stimulation of median raphe terminals in dorsal CA2 reduces social investigation in male mice specifically investigating social stimulus of ovariectomized female mice

**DOI:** 10.1101/2023.08.30.555504

**Authors:** Su Hyun Lee, Nicholas I. Cilz, Sarah K. Williams Avram, Adi Cymerblit-Sabba, June Song, Karis Courey, Austin Howley, Michaela E. Cooke, W. Scott Young

**Affiliations:** Section on Neural Gene Expression, National Institute of Mental Health, National Institute of Health, Bethesda, MD, United States

**Keywords:** Social behavior, social interaction, CA2, serotonin, median raphe nucleus, optogenetics

## Abstract

The cornu ammonis area 2 (CA2) region is essential for social behaviors, especially in social aggression and social memory. Recently, we showed that targeted CA2 stimulation of vasopressin presynaptic fibers from the paraventricular nuclei of hypothalamus strongly enhances social memory in mice. In addition, the CA2 area of the mouse hippocampus receives neuronal inputs from other regions including the septal nuclei, the diagonal bands of Broca, supramammillary nuclei, and median raphe nucleus. However, the functions of these projections have been scarcely investigated. A functional role of median raphe (MR) – CA2 projection is supported by the MR to CA2 projections and 82% reduction of hippocampal serotonin (5-HT) levels following MR lesions. Thus, we investigated the behavioral role of presynaptic fibers from the median raphe nucleus projecting to the dorsal CA2 (dCA2). Here, we demonstrate the optogenetic stimulation of 5-HT projections to dCA2 from the MR do not alter social memory, but instead reduce social interaction. We show that optical stimulation of MR fibers excites interneurons in the stratum radiatum (SR) and stratum lacunosum moleculare (SLM) of CA2 region. Consistent with these observations, we show that bath application of 5-HT increases spontaneous GABA release onto CA2 pyramidal neurons and excites presumed interneurons located in the SR/SLM. This is the first study, to our knowledge, which investigates the direct effect of 5-HT release from terminals onto dCA2 neurons on social behaviors. This highlights the different roles for these inputs (i.e., vasopressin inputs regulating social memory versus serotonin inputs regulating social interaction).

## Background

The ability to recognize a conspecific is essential for survival, given its importance for reproduction, pair-bonding, and parental behaviors. The cornu ammonis area 2 (CA2) region is essential for social behaviors and is implicated in social aggression and social memory (1-7). In patients with schizophrenia and bipolar disorder, syndromes that exhibit deficits in social interactions, the numbers of non-pyramidal cells in the specifics of CA2 region are reduced (8). Recent studies showed that genetically targeted inactivation of CA2 pyramidal neurons causes a deficit in social memory (6), and stimulating vasopressin presynaptic fibers projecting from the paraventricular nucleus of the hypothalamus to the dorsal CA2 (dCA2) enhances social memory in mice (3). Other extrahippocampal inputs to dCA2 include the median raphe nucleus, medial septum, and supramamillary nucleus (9). However, the behavioral roles of these inputs to dCA2 are relatively unknown.

Serotonin (5-HT) plays important roles in various behaviors including aggression, anxiety, depression, and parental care (10-13). Hyperserotonemia, or elevated blood 5-HT, is found in 25-35% of patients with autism spectrum disorder (14, 15). This elevation may be due to the increased 5-HT synthesis capacity found in autistic children, as it was shown that 5-HT synthesis capacity was higher in older autistic children and men compared to controls (16). In rats, administration of the selective serotonin reuptake inhibitor (SSRI), citalopram, reduced social interaction, a trait of autism, through enhanced activation of the 5-HT2c receptor (17).

Within the brain, serotonergic neurons are found in bundles that project caudally to the brain stem and spinal cord or rostrally to the forebrain. The rostral group contains approximately 85% of the 5-HT neurons and includes the interpenduncular, caudal linear, dorsal and median raphe (MR) nuclei (18). In mice, Cui *et al*. have shown that most, but not all, serotonergic dCA2 projections arise from the MR nucleus (9). In CA1 and CA3, it was previously shown that interneurons throughout the stratum radiatum (SR) and oriens of the hippocampus receive inputs from MR (19) and optogenetic stimulation of MR fibers in the hippocampus directly excites interneurons via activation of ionotropic 5-HT3 and glutamatergic receptors in CA1 (20). However, single pulse photostimulation did not evoke detectable responses in CA3 and CA1 pyramidal cells (20). The direct effects of evoked 5-HT release from serotonergic input to the hippocampus, specifically to the dCA2, have not been investigated.

Based on these previous studies, we hypothesized that if we optogenetically stimulated in the dCA2 the serotonergic fibers from the MR nucleus, we would regulate social behavior. We ran a battery of behavioral tests involved in social, anxiety-like, repetitive and compulsive-like, and locomotor activities. We found that stimulating serotonergic presynaptic fibers from the MR nucleus projecting to dCA2 did not alter social recognition memory (SRM), unlike stimulating vasopressin fibers in dCA2 originating from paraventricular nucleus (3). However, unexpectedly, this stimulus paradigm did decrease social interaction in male mice with social stimulus of ovariectomized (OVX) female mice when presented with both a social as well as an inmate object, suggesting a projection involved in the regulation of social interaction in male mice.

## Materials and Methods

### Subjects

All experiments were conducted according to US National Institutes of Health guidelines for animal research and were approved by the National Institute of Mental Health Animal Care and Use Committee. Adult male 5-HT transporter promoter (Slc6a4 gene)-driven Cre recombinase (Cre) [Tg(Slc6a4-cre) ET33Gsat mice, generously provided by Charles Gefen, NIMH] (Raphe-cre) mice were back-crossed for more than 10 generations into the C57BL/6J strain (Jackson Laboratory, Bar Harbor, ME, USA). The 5-HT transporter is expressed selectively in 5HT neurons (2, 21). All mice were singly housed during experiments and maintained on a 12-h light cycle (lights off at 1500 h) with ad libitum access to food and water. We chose to singly house the mice as described in Smith *et al*. as to follow our in-house standard protocol where they do not remember conspecific after the two-hour interval for the social recognition memory test. Power analysis was not performed prior to the experiment as there was no prior study looking at serotonergic transporter manipulation in dCA2 to our knowledge.

For the tracing studies, the Raphe-cre line was used for anterograde tracing. Furthermore, the Raphe-cre line was crossed with B6.129(Cg)-Gt(ROSA)26^*sortm4luo/J*^ (22) to express green fluorescent protein (GFP) in 5-HT neurons. BalbC female mice (Jackson Laboratory) were ovariectomized in-house. OVX BalbC female mice were used as stimulus choices for behavioral testing as based on extensive pilot work in our lab showing the strongest and most consistent response from male C57Bl/6J mice. Stimulus mice were used only once per day. Please refer to supplementary methodology for the detailed procedure.

For electrophysiology studies using bath application of 5-HT, offspring of Avpr1b-Cre mice (23) crossed with the Ai9 td-Tomato expressing reporter line (Jackson Laboratory) were used to identify the CA2 region. Please refer to supplementary methodology for detailed information.

### Stereotaxic surgery, viral tracing, and optogenetic surgical implantation

For anterograde viral tracing, Raphe-cre mice were injected with Cre-dependent rAAV2-CAG-FLEX-EGFP (500 nL of 2 × 10^+,^ transducing units per ml), serotype 2, in the MR (ML:0.00 mm, AP: -4.2 mm, DV:-4.5 mm) and incubated for 4 weeks. For optogenetic behavioral studies, Cre-dependent AAV-DIO-EF1α-hChR2(H134R)-mCherry (500 nL of 1.5 × 10^+,^ transducing units per ml), serotype 2, was injected into the MR nucleus. 200 um optic fiber was implanted in dCA2 region. Please refer to supplemental methodology section for detailed methodology of stereotaxic surgery, viral tracing and optogenetic surgical implantation.

### Histology and immunohistochemistry

Brains were processed with immunohistochemistry to detect cells expressing 5-HT and GFP in tissues. See supplemental methods for detailed procedure of histology and immunohistochemistry.

### Electrophysiology

Please refer to supplemental method section for detailed methodology of whole-cell recordings obtained from Raphe-Cre mice after 5-12week viral infection and bath 5-HT recordings from slices prepared from offspring of Avpr1b-Cre mice crossed with an Ai9 reporter line.

### Behavioral tests

Behavioral tests were conducted 6 to 8 weeks after surgery involving viral injection followed by probe implantation. All behavioral tests were conducted during light-cycle between 9:00 AM and 2:00 PM. During the behavioral tests with photostimulation, ChR2-positive fibers were stimulated by a LED light source (465 nm, PlexonBright; and LED driver LD-1; Plexon, Dallas, TX) with 20-ms pulses at 20 Hz using a programmable pulse stimulator (NI USB-6229; National Instrument, Austin, TX) and LabView software (version 12.0.1f4; National Instruments), with a power output of 1-2 mW/mm^2^. Mice were habituated to the testing room for 30 mins prior to behavioral testing and optogenetic cables for 5 mins prior to all the behavioral tests except open-field and marble burying tests as these are a measure of anxiety-like behavior. Please refer to supplemental methods section for the behavioral measurements of SRM test, object recognition, social interaction, and open-field tests.

### Social recognition memory

SRM tests were conducted in a new mouse cage during the light cycle. Behaviors were recorded using Ethovision XT (version 13; Noldus Information Technology, Leesburg, VA). During acquisition trials, mice were exposed to a freely moving unfamiliar OVX mouse in a cage for 5 min. After 30-min and 2-hour retention intervals in the absence of the stimulus female, retrieval trials were performed in which mice were exposed to the same familiar OVX female for 5 min. For 2-hour retention intervals, mice were photostimulated during the acquisition trial as this was how memory effects were studied previously (3).

### Object recognition

Object recognition acquisition was conducted in the same manner as the SRM tests, except that a scintillation vial filled with purple solution was used instead of a female mouse.

### Social interaction test

Social interaction tests were conducted during the light cycle in a three-chamber apparatus made out of clear plexiglass (width: 27.5 cm x length: 56 cm x height: 23 cm and gate opening of 11 cm) with the scintillation vial described above and a tethered OVX female in opposite outer chambers. The OVX female mice for 30 mins prior to testing were habituated to a collar and metal leash fixed to the wall of the chamber allowing direct contact of test mice with the OVX female mice. The trials were conducted for 10 min and mice were photostimulated during the trial. Tests were recorded using Ethovision XT.

### Open-field test

Mice were placed in an opaque white open-field box (35 cm × 35 cm × 35 cm) for 10 min during optogenetic stimulation. Locomotor activity and anxiety-like behavior (degree of thigmotaxis) were measured. Mice were tracked with Ethovision XT software.

### Marble burying test

Twelve marbles were placed on top of the bedding spread across the mouse cage (4 by 3) on a layer of woodchip bedding 5 cm deep. The mice were optogenetically stimulated across 30 mins and the latency to dig was measured and the number of marbles buried was counted.

### Statistical tests

Two-way repeated measures of ANOVA (interaction of genotype x trial) and post-hoc Sidak’s multiple comparison tests were performed, using the Prism 7.0 software (GraphPad, La Jolla, CA, USA). Normality was checked using Shapiro-Wilk normality test. Ordinary one-way ANOVA was used to compare between genotypes and post-hoc Turkey’s multiple comparison test was performed subsequently.

## Results

### Projections from MR to the dCA2

Cui *et al*. (9) previously showed that there are subcortical projections to the dCA2 from MR in the mouse. Here, we validated those projections from the MR by using viral tracing and transgenic approaches. After injecting AAV-CAG-FLEX-EGFP in the MR of Raphe-cre mice (Fig. 1A-B), GFP expression was prominent in both MR neurons (Fig. 1B-D) and fibers projecting to the dCA2 (Fig. 1E). Immunostaining for 5-HT (Fig. 1C) confirms GFP expression is restricted to 5-HT neurons in MR. The transgenic Raphe-cre line crossed with the GFP reporter line shows GFP expression in MR neurons and fibers projecting to the dCA2 (Fig. 1F-G).

**Figure 1.**
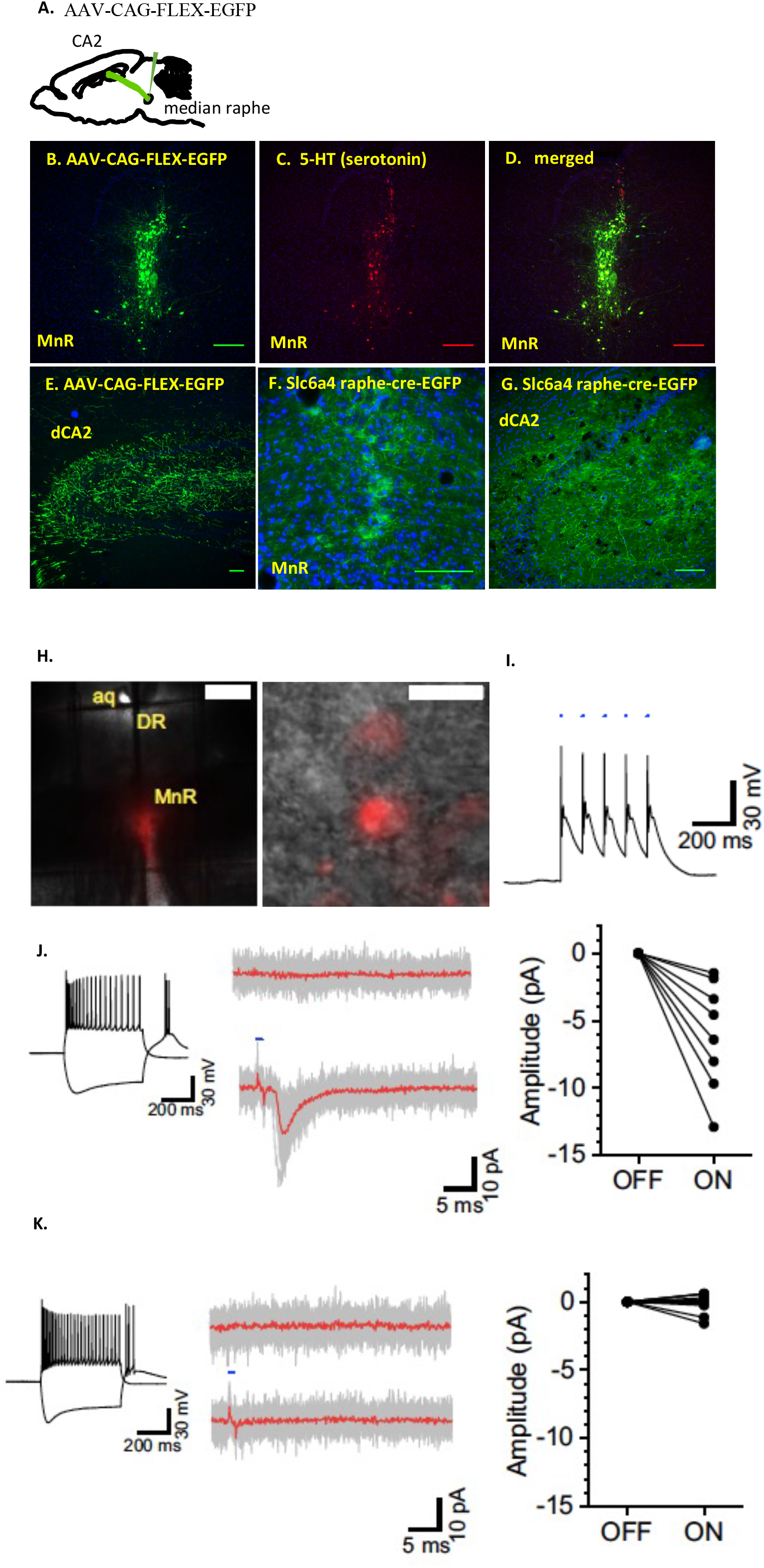
Serotonin neurons from the median raphe project to the dorsal CA2. A. Diagram illustrating, for panels B-E, a Raphe-cre mouse injected with AAV-CAG-FLEX-EGFP in the MR region and fibers projecting to dCA2. B. Resulting GFP (green) expression in MR neurons. C. 5-HT immunohistochemical staining of 5-HT (red) in the MR region. D. Merged image of 5-HT (red) and GFP (green) with colocalization (yellow). E. MR fibers projecting to dCA2 from the MR. F. Transgenic Slc6a4 Raphe-cre mouse crossed with B6.129(Cg)-Gt(ROSA)26^Sortm4(ACTB- tdTomato,-EGFP)Luo^/J mouse shows GFP expression in MR neurons and in panel G fibers projecting to dCA2. Scale bars for panels B-G are 50 μm. H. *Left*, Representative coronal slice of MR region displaying robust mCherry reporter expression of ChR2 virus. *Right*, Representative image of transduced cells in MR. Scale bars are 200 and 10 µm, respectively. I. Representative light-induced action potential firing of MR neurons. J. *Left*, Input/output response of light-responsive interneuron patched in the SR region proximal to CA2. *Middle*, Light-induced excitatory postsynaptic current elicited from the same neuron. *Right*, Summary data of normalized amplitudes seen with light stimulation. K. *Left and middle*, Input/output response of a non-light-responsive interneuron patched in the SR region proximal to CA2. *Right*, Summary data of normalized amplitudes seen with light stimulation for non-responsive cells.

### dCA2 interneurons located in the stratum radiatum and stratum lacunosum moleculare receive input from the median raphe

We verified the functionality of our ChR2 injections by first making whole-cell recordings in slices prepared from MR transduced Raphe-Cre mice. Current-clamp recordings from the MR neurons demonstrate that brief (2 ms) pulses of light evoked reliable action potential firing in mCherry-positive neurons (Fig. 1H-I). Hippocampal interneurons receive input from the median raphe (19) and optogenetic stimulation of MR fibers in the hippocampus directly excites interneurons via activation of ionotropic 5-HT3 and glutamatergic receptors (20). We therefore used light-evoked excitatory postsynaptic currents (EPSCs) as a proxy to indirectly assess the functionality of MR terminal activation in the dCA2 by recording from presumed interneurons located in the SR and stratum lacunosum moleculare (SLM) of the CA2 region. We observed small but reliably evoked EPSCs in 8 of 25 cells. We also observed 1 cell that showed a pronounced membrane hyperpolarization in response 10 stimuli delivered at 20 Hz (Supplemental Fig. 2). Overall, 34.6% of cells in the dCA2 responded to optogenetic stimulation (Fig. 1J-K).

### Optical stimulation of MR fibers within the dCA2 reduces social interaction but not recognition

Virus expressing Cre recombinase-activatable channelrhodopsin fused with mCherry was injected into the MR of both Raphe-cre and WT mice (Supplemental Fig. 1A). mCherry expression was observed in the cell bodies of the MR and fibers within the dCA2 region from the former strain indicating expression and transport of channelrhodopsin from the MR to the dCA2 (Supplemental Fig. 1B and C). Bilateral optic fiber implants were placed just above the dCA2 and the terminals within the CA2 region for blue light stimulation (Supplemental Fig. 1D). We conducted two-trial SRM tests using ovariectomized (OVX) stimulus mice with or without optical stimulation. The interval between the trials was either 30 min or two hours during which male WT mice either remember familiar mice or not, respectively (Fig. 2A-D and 2G-I,). In the 30-min interval trials there were (A) Raphe-cre-ON-OFF group: Raphe-cre group optogenetically stimulated only during acquisition period. (B) Raphe-cre-ON-ON group: Raphe-cre group optogenetically stimulated during acquisition and retrieval period. (C) Raphe-cre-OFF-OFF group: Non-stimulated Raphe-cre group. (D) WT-ON-ON group: WT group optogenetically stimulated during acquisition and retrieval period (Fig. 2A-D). Non-stimulated Raphe-cre mice and WT mice sniffed significantly less during retrieval period when compared to acquisition period (Raphe-cre-OFF-OFF, *p* = 0.0005; WT-ON-ON, *p* = 0.01) (Fig. 2E). When stimulated during only acquisition or both acquisition and retrieval periods, there were no significant differences in sniffing between the two periods (Raphe-cre-ON-OFF, NS, *p* = 0.99; Raphe-cre-ON-ON, NS, *p* = 0.011) (Fig. 2E). A decrease in the sniffing ratio (calculated by Retrieval sniffing duration - Acquisition sniffing duration/Acquisition sniffing duration^x10^0) between acquisition to retrieval trials of >30% indicates that the test mouse remembers the OVX stimulus mouse. Raphe-cre mice stimulated during the acquisition (ON-OFF) period only had significantly higher sniffing ratios (*i*.*e*., absence of sniffing changes) compared to all other groups (Raphe-cre ON-OFF compared to Raphe-cre ON-ON, Raphe-cre OFF-OFF, and WT ON-ON; *, *p* <0.05; ##, *p* <0.01; and +, *p* <0.05 respectively) suggesting that lack of change in ratio was driven by decreased sniffing investigation in the acquisition trial (Fig. 2F). The significant decrease in sniffing ratios for the Raphe-cre-ON-ON group when compared to the Raphe-cre-ON-OFF group suggests that SRM was present for this Raphe-cre optogenetically stimulated group (*p* <0.05) (Fig. 2F). We further conducted 2-trial SRM memory tests with 2-hour intervals (Fig. 2G-I) and neither Raphe-cre nor WT mice showed SRM (Fig. 2J - K), whether stimulated or not during acquisition. This indicates that dCA2 stimulation of the 5-HT fibers did not stimulate social memory. In order to emphasize that we are observing an effect of stimulation during acquisition on social interaction, not memory, which accounts for the significantly higher sniffing ratio of the Raphe-cre-ON-OFF group in 30-min interval trials, we compared sniffing durations obtained during all the optogenetic stimulations during the acquisition period. Overall, there was significant decrease in sniffing in Raphe-cre mice compared to WT mice when optogenetically stimulated during the acquisition period (Raphe-cre stimulation-ON vs WT-cre stimulation-ON, p<0.004) (Fig. 2L).

**Figure 2.**
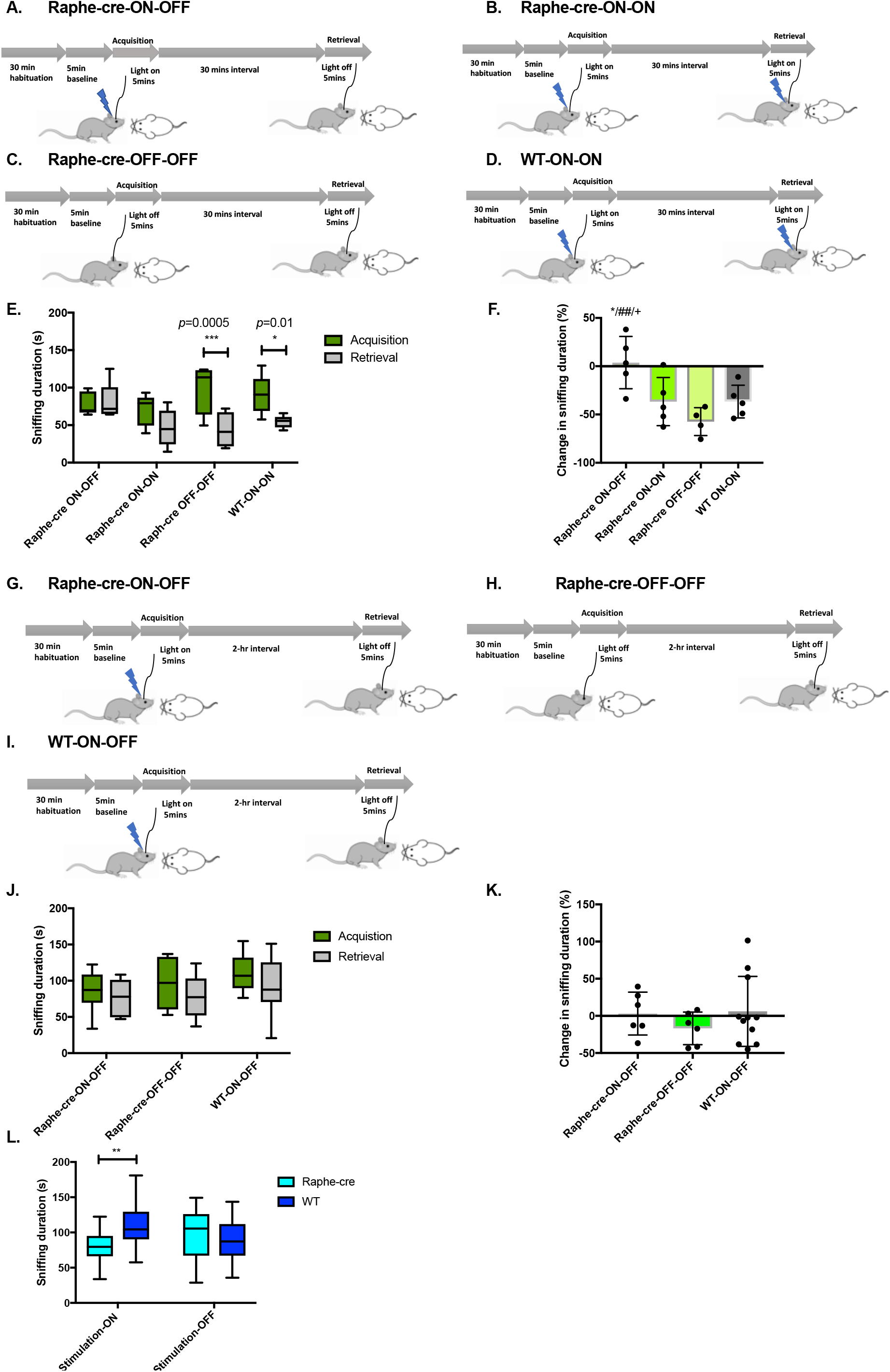
Social recognition memory is unaffected while social interaction is reduced. A-F. The subject interacts with an ovariectomized (OVX) Balb/C female for 5 minutes and after a 30-minute interval, is exposed to the same mouse (familiar test). A. Raphe-cre group optogenetically stimulated during acquisition period (n = 5). B. Raphe-cre group optogenetically stimulated during acquisition and retrieval period (n = 5). C. Non-stimulated Raphe-cre group (n = 4). D. WT group optogenetically stimulated during acquisition and retrieval period (n = 5). Animals without viral expression, mis-targeting of fiber optics in stimulated Raphe-cre and WT group were excluded from the analysis. E. Sniffing duration during acquisition and retrieval trial. Non-stimulated Raphe-cre mice and WT mice sniffed significantly less during retrieval period when compared to acquisition period (two-way repeated measures of ANOVA: genotype x trial F(3,15)=5.4 *p* <0.01, post-hoc Sidak’s multiple comparison test Raphe-cre-OFF-OFF, ***, *p* = 0.0005; WT-ON-ON, *, *p* = 0.01). When stimulated during only acquisition or both acquisition and retrieval periods, there were no significant differences between the acquisition and retrieval periods (post-hoc Sidak’s multiple comparison test Raphe-cre-ON-OFF, NS, *p* = 0.99; Raphe-cre-ON-ON, NS, *p* = 0.11). F. Change in sniffing duration calculated by (Retrieval sniffing duration − Acquisition sniffing duration)/Acquisition sniffing duration^x^ 100. Raphe-cre mice stimulated during the acquisition period had significantly higher sniffing ratios compared to all other groups (one-way ANOVA F(3,15)=6.3 *p* <0.005, post-hoc Turkeys multiple comparison test shows that change in sniffing ratio of Raphe-cre ON-OFF group was significantly different to Raphe-cre ON-ON, Raphe-cre OFF-OFF and WT ON-ON, *, *p* <0.05; ##, *p* <0.01; and +, *p* <0.05 respectively). G-K. SRM test with 2-hour interval. G. Raphe-cre group optogenetically stimulated acquisition period (n = 6). H. Non-stimulated Raphe-cre group (n = 6). I. WT group optogenetically stimulated acquisition period (n = 11). Animals without viral expression, mis-targeting of fiber optics in stimulated Raphe-cre and WT group were excluded from the analysis. J. Sniffing duration during acquisition and retrieval trial. Sniffing duration during acquisition and retrieval period were not different across genotypes and stimulations. K. Change in sniffing duration calculated as above. There was no significant difference between the genotypes and stimulations. L. Sniffing duration of OVX female mice during optogenetic stimulation. Groups include Raphe-cre group optogenetically stimulated (n = 18), WT group optogenetically stimulated (n = 21), Non-stimulated Raphe-cre group (n = 18), and Non-stimulated WT group (n = 16). Two-way ANOVA: genotype x trial F(1,69)=8.04, *p* <0.006. Post-hoc Šídák’s multiple comparison test were performed between the genotypes, the sniffing duration for Raphe-cre group was significantly lower when optogenetically stimulated. Raphe-cre stimulation-ON vs WT stimulation-ON light on session. **, *p* <0.004.

We conducted three-chamber social interaction tests in order to determine whether stimulation of the 5-HT presynaptic fibers within the dCA2 alters the preference for social over novel object interaction (Fig. 3A). Mice are placed in the middle of the chamber and exposed to an inanimate object in one outer chamber and a novel ovariectomized Balb/C female in the other outer chamber for 10 minutes during light stimulation. As is the case with the 2-trial SRM test above, the stimulated Raphe-cre mice sniffed the OVX female mice significantly less in comparison to WT mice (Fig. 3B, the main effect is driven by the interaction between the genotype and OVX female being explored, *p* <0.001). There was no significant difference between groups when not stimulated (Fig. 3C). Furthermore, the sniffing ratio of mouse over total sniffing was significantly lower for Raphe-cre mice when light-stimulated (Fig. 3D, *p* <0.01), while there was no significant difference when not stimulated (Fig. 3E). The sniffing durations for object investigation were low and not affected by light stimulation (Fig. 3B and 3C).

**Figure 3.**
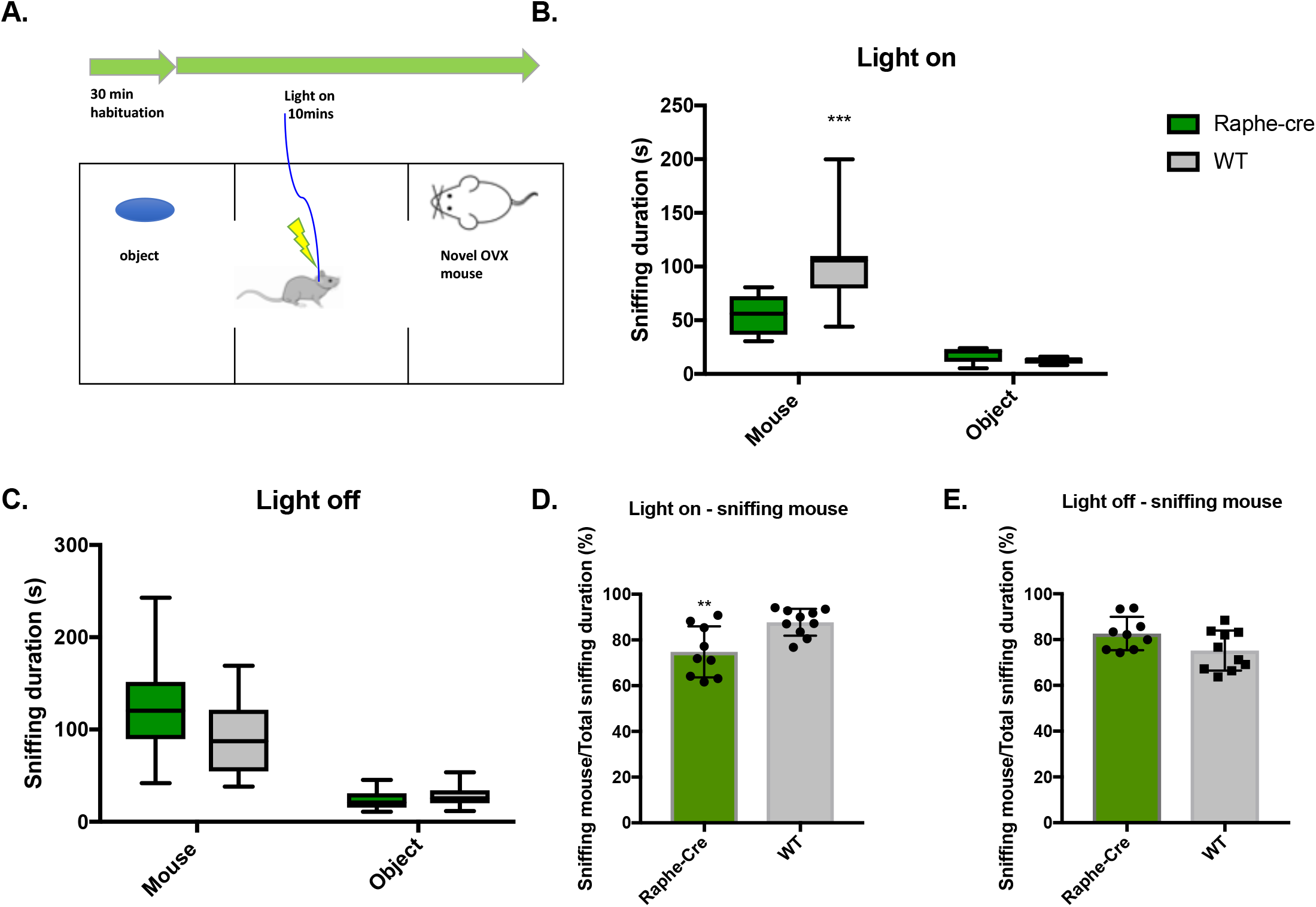
A. Social interaction is reduced in the three-chamber apparatus. B. Sniffing duration was significantly different between the genotype of Raphe-cre (n = 9) and WT (n = 10) and this effect was driven by the sniffing of OVX female mice. Animals without viral expression, mis-targeting of fiber optics in stimulated Raphe-cre and WT group were excluded from the analysis. One-way Repeated Measures ANOVA: genotype x the sniffing of OVX female and object being explored (F(1, 17)=11.33, *p*<0.01). Main effect of genotype (F(1, 17) = 8.82, *p* = 0.01) and main effect between the sniffing of OVX female and object being explored (F(1,17) = 61.87, *p* <0.001) was found. The main effect is driven by the interaction between the genotype and OVX female being explored using the post-hoc Šídák’s multiple comparison test (***, *p*<0.001). One outlier was identified and removed in WT group sniffing the object using Grubb’s test and normal distribution was maintained. C. No optogenetic stimulation. D. Sniffing mouse/total sniffing duration (%) with light on (unpaired t-test, *, *p* <0.01). E. Sniffing mouse/total sniffing duration (%) no optogenetic stimulation.

### Light stimulation of MR fibers within the dCA2 does not alter object recognition

Object recognition was conducted in a new mouse cage in which a 100 mL scintillation vial filled with purple solution was used as an object and placed in the cage for 5 min. Mice were re-exposed to the same familiar object after a 2-hour retention interval. Stimulation of the presynaptic 5-HT fibers in the dCA2 did not enhance or inhibit object memory or interaction (Supplemental Fig. 3A-E).

### Optical stimulation of MR fibers within the dCA2 does not alter anxiety-like behaviors or locomotor activity

Twelve marbles were placed in the mouse cage with 5 cm of woodchip bedding and mice were optically stimulated. The latency to dig was measured for 5 minutes and the number of marbles buried after 30 minutes was counted (Fig. 4A). 5-HT terminals stimulation within dCA2 did not effect the mice latency to dig nor the number of buried marbles (Fig. 4B – C). Mice were placed in an opaque white open-field box for 10 min during light stimulation (Fig. 4D). As controls, these mice went through open-field testing without light stimulation on a different day. There was no significant difference in locomotor activity between Raphe-cre and WT mice (Fig. 4E). There were no significant differences in center duration or frequency to enter the center zone between Raphe-cre and WT mice when stimulated suggesting no effect on anxiety-like behaviors (Fig. 4F and G).

**Figure 4.**
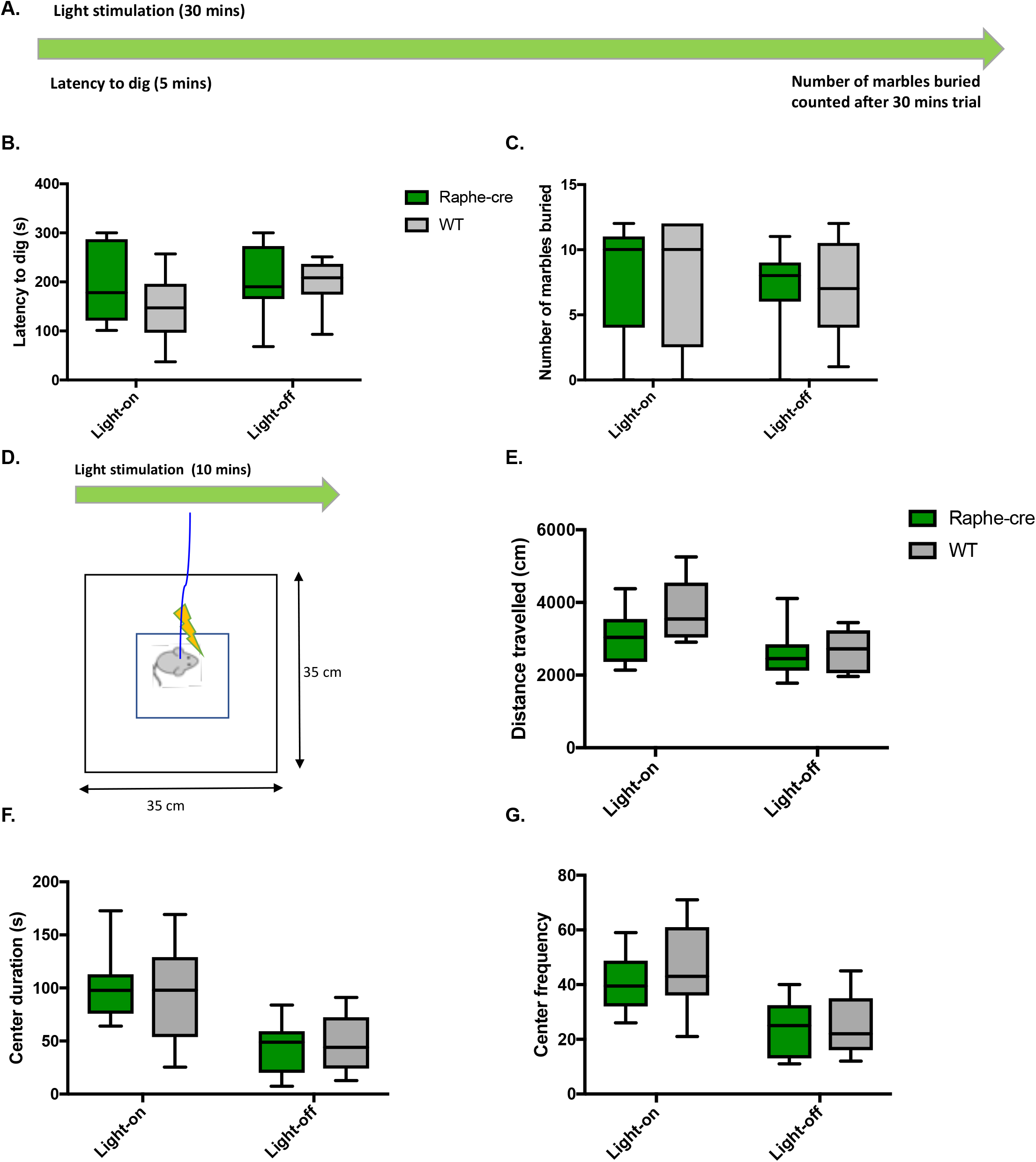
Anxiety-like Behaviors are unaffected. A. 30-min marble burying test. B. Latency to dig while light-on (Raphe-cre: n = 7; WT: n = 13) and off (Raphe-cre: n = 7; WT: n = 6). C. Number of marbles buried while light-on and off. D-G. 10-min open field test. D. Illustration of open-field test setup. E. Total distance travelled in open-field box light-on (Raphe-cre: n = 8; WT: n = 11) and off (Raphe-cre: n = 13; WT: n = 11). F. Center distance travelled. G. Center frequency. Animals without viral expression or mis-targeting of fiber optics in the stimulated Raphe-cre group were excluded from the analysis.

### 5-HT augments GABAergic transmission by exciting CA2 hippocampal interneurons

The modulatory effects of 5-HT on the CA2 region remain unclear. Since we show that stimulating MR fibers in the CA2 region directly increases interneuron excitability (See Fig. 1) and because 5-HT augments GABA release elsewhere in the CA1 region (24, 25), we next examined whether 5-HT alters inhibitory drive onto CA2 neurons to begin to explore the underlying cellular mechanisms of 5-HT in the CA2 region. To selectively study the CA2 region in vitro, we made slices from offspring of Avpr1b-Cre mice (23) crossed with the Ai9 td-Tomato expressing reporter line (Fig. 5A and B) (26) and made whole-cell recordings of spontaneous inhibitory postsynaptic currents (sIPSCs) from Avpr1b-expressing neurons. Bath application of 5-HT (30 µM) significantly increases the frequency of sIPSCs onto CA2 neurons (Control: 2.45±0.4 Hz, 5-HT: 7.92±1.6 Hz, n = 6, *P* = 0.008, paired sample t-test, Fig. 5C, D, and E). Following washout of 5-HT, subsequent application of the selective GABA^A^ antagonist bicuculline (20 µM) abolishes sIPSC events, indicating that they are GABAergic (Fig. 5C and D). To determine whether action potential generation is necessary for 5-HT-induced increases in sIPSCs, we included additional experiments with TTX (0.2 μM) in the recording solution to measure miniature inhibitory postsynaptic currents (mIPSCs). Under these conditions, 5-HT fails to increase mIPSC frequency (Control: 1.65±0.3 Hz, 5-HT: 1.45±0.3 Hz, n = 4, Figure 5F). The requirement of action potential generation in 5-HT-induced increases in sIPSCs suggests that 5-HT may act on presynaptic interneurons to increase their excitability. We therefore made whole-cell current-clamp recordings from interneurons proximal to the CA2 region (Fig. 5G). Bath application of 5-HT depolarizes interneurons by ∼5 mV (Control: -63.4±1.3 mV, 5-HT: -56.3±2.3 mV, n = 6, P = 0.02, paired sample t-test, Fig. 5H and I).

**Figure 5.**
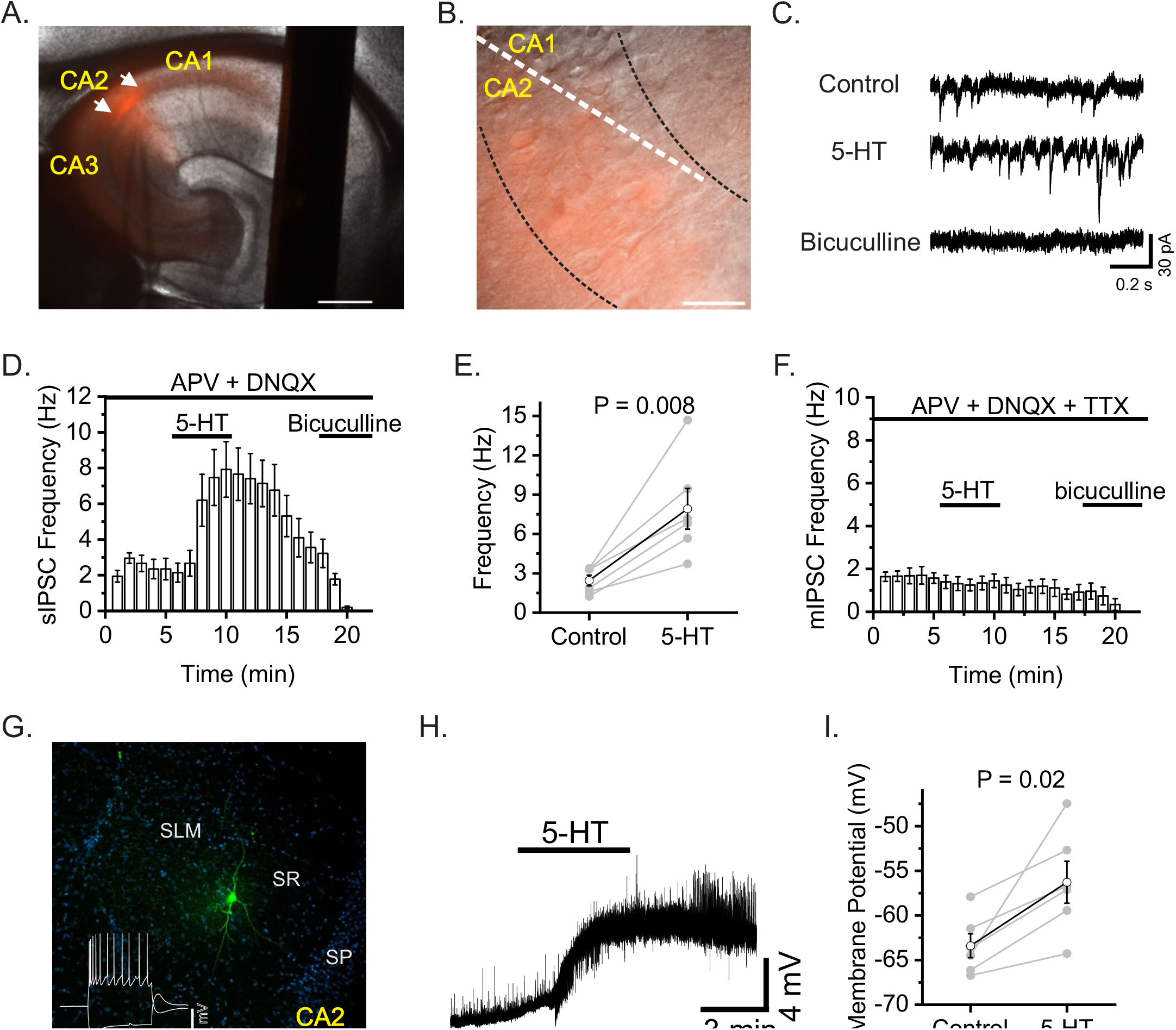
5-HT increases GABA release onto CA2 pyramidal neurons. A and B. Representative hippocampal section of Avpr1b-Cre x Ai9 showing td-Tomato positive neurons. C. sIPSC traces made from CA2 neuron before 5-HT, during 5-HT, and during bath application of bicuculline. D. Time course of sIPSCs recorded from CA2 neurons (n=6). E. Summary data of sIPSC frequency for 5 min control versus last minute of 5-HT. F. Time course of mIPSCs recorded from CA2 neurons (n=4). G. Biocytin labeling of a typical SR interneuron proximal to CA2. H. Representative trace illustrating the depolarizing effect of 5-HT on CA2 proximal interneurons. I. Summary data for interneuron membrane potential before and after application of 5-HT (n=6).

## Discussion

The dCA2 region is essential for social behaviors (1-7), especially in social aggression and social memory. A functional role of the MR - CA2 microcircuit is suggested by the MR to CA2 projections (9), the presence of hippocampal interneuron input from the MR (19), and the 82% reduction of hippocampal 5-HT levels following MR lesions (27). Here, we demonstrate that optogenetic stimulation of 5-HT projections from the MR does not alter social memory, but instead reduces social interaction with OVX female mice. This is the first study, to our knowledge, that investigates the direct effect of 5-HT release from terminals onto dCA2 neurons on social behaviors. We also show that stimulating MR fibers in the SR and SLM of CA2 region excites interneurons. Furthermore, we explored mechanisms of 5-HT in the CA2 region and showed 5-HT augments inhibitory drive onto CA2 neurons by depolarizing presumed interneurons near the CA2 region.

In this study, our transgenic and various viral tracing approaches confirmed afferent projections from the MR to dCA2 that we previously observed using retrograde beads that labeled most but not all of the MR nucleus (9). Coronal slices containing the MR region displayed mCherry reporter expression from the ChR2 virus. After confirming the axonal projection to the dCA2 from the MR, we conducted an initial pilot study of two-trial SRM test. The interval between the trials were 30-min and 2-hours, the shorter interval permitting recall with the longer one not, allowing for the study of memory enhancement. Overall, optogenetically stimulating these projections did not enhance nor inhibit the SRM. Therefore, we did not invest SRM further. Although, our initial study showed that optogenetically stimulating projections from the MR to the dCA2 during acquisition period (Raphe-cre-ON-OFF) had significantly higher sniffing ratio in 30-min interval trials, it was not due to SRM memory deficit but rather a decrease in sociability. There was a significant decrease in sniffing duration in Raphe-cre mice compared to WT mice when these mice were optogenetically stimulated during the acquisition period suggesting that projections from the MR to the dCA2 play a role in modulating sociability. Furthermore, to validate the effect of decrease in sociability in freely moving mice as described above, we conducted full trial of three-chamber social interaction tests to determine whether stimulation of the 5-HT presynaptic fibers within the dCA2 alters the preference for social over novel object interaction. The stimulated Raphe-cre mice sniffed the OVX female mice significantly less in comparison to WT mice. Our results also show that the role of fibers projecting from the MR to the dCA2 is different from the vasopressin fibers projecting from paraventricular nucleus to the dCA2 (3).

Serotonin’s key role in shaping social responses is well recognized. Studies describing the role of 5-HT and sociability across the evolutionary spectrum have included enhanced prosocial behaviors in the octopus after the administration of methylendioxymethamphetamine (MDMA), which principally binds to the site of serotonin transporter (SERT) (28, 29). In non-human primates, 5-HT plays an important role in directing attention to monitoring other individuals (30).

Walsh *et al*. (29) demonstrated that stimulating 5-HT release in the nucleus accumbent (NAc) promotes sociability through activation of NAc 5-HT1b receptors. Furthermore, inhibition of dorsal raphe (DR) 5-HT activity or their terminals in the NAc reduces social interaction (29). However, it appears that the anatomical origins and destinations of 5HT innervation differently affect social behaviors. In contrast to enhanced social behaviors just described above, in this study we show a decrease in sociability when MR fibers in the CA2 region are stimulated. The dCA2 is richly innervated by 5HT fibers from the MR (Fig. 1 and Cui *et al*. (9)). The specificity of anatomical differences between MR nucleus and DR may play a different role in social behavior. Earlier studies have demonstrated that lesions in DR produced a non-significant 10 % reduction in hippocampal 5-HT level; on the other hand, lesions in MR lead to 82 % reduction in 5-HT level (27).

We show that stimulating MR fibers in the dCA2 region increases interneuron excitability, and it was previously shown that optogenetic stimulation of MR fibers in the hippocampus directly excites interneurons via activation of ionotropic 5-HT3 and glutamatergic receptors (20). Furthermore, we demonstrate that 5-HT alters inhibitory drive onto dCA2 neurons by increasing GABA release onto dCA2 pyramidal neurons. In the future, validation of agonist bath application sIPSC experiments with an optogenetic approach and examination of which 5-HT receptors are involved in augmenting GABAergic transmission are necessary. Moreover, for the future experiments, it would be interesting to simultaneously demonstrate whether the effect is mediate by 5-HT and not glutamate or combination of the two involving pharmacological applications. Furthermore, a recent hippocampus RNA-seq study showed that 5-HT7 receptors are highly expressed in the dCA2 pyramidal cells to at a least 5-fold greater level than in the dCA1, CA3, and dentate gyrus neurons (31). Thus, it would be interesting to explore the role of 5-HT7 receptors in dCA2 pyramidal cells.

In our study, optogenetic stimulation of Raphe-cre fibers projecting to the dCA2 did not alter object recognition memory. In a recent study, oxytocin receptors in anterior CA2/CA3 and dentate gyrus were shown to be needed for social discrimination, but not the non-social cues (7). Furthermore, stimulating vasopressinergic fibers to the dCA2 did not enhance object memory (3). Thus, these studies agree that CA2 does not play a major role in object memory.

Earlier studies have originally proposed marble burying as an animal model of anxiety in which a decrease in marble burying was seen with intraperitoneal injection of serotonin reuptake inhibitors such as citalopram and benzodiazepines (32). In the current study, optogenetic stimulation of Raphe-cre terminals within the dCA2 did not alter latency to dig or number of marbles buried. More recently, marble burying adequately conceptualizes more of a preclinical animal model of compulsive or repetitive behavior (32, 33). Thus, our results suggest that the projection from the MR to the dCA2 is not involved in this role of 5-HT. Furthermore, we did not observed anxiety-like behavior based on time spent in the center of an open-field arena, nor locomotor activity. This indicates that the decrease in social interaction was not due to an alteration in anxiety-like behavior. Furthermore, although we chose marble burying as we were testing specifically effect of serotonergic circuits, anxiety-like behavior can be further validated using O-maze or Elevated Plus Maze system.

To our knowledge, this is the first study that has investigated the role of stimulation of presynaptic fibers projecting from the MR to dCA2 in social behavior. In conclusion, we highlight that this manipulation of inputs to the CA2 did not alter the SRM but did decrease social interaction in male mice with OVX female mice. We show that stimulating MR fibers in the CA2 region directly excites interneurons and that 5-HT application increases inhibitory drive onto CA2 neurons by depolarizing presumed interneurons. We observed no effects on social or object memory, or anxiety-like behavior. Interestingly, increasing inhibitory drive onto dCA2 neurons results in a different effect than substantial inhibition of dCA2 pyramidal neurons themselves. This suggests the region may be involved in other more subtle social environment monitoring that can be tested with more specific behavioral protocols. Future studies will be necessary to develop insight into how this action translates to the understanding of neuropsychiatric disease.

## Supporting information

NIH Publishing Agreement & Manuscript Cover Sheet

## Acknowledgement

A special thank you to everyone in the Section on Neural Gene Expression for their contributions and hard work. We would like to thank Shweta Sahu and Ruby Kang for their assistance during stages of data collection. This research was supported by the National Institute of Mental Health (NIMH) Intramural Research Program (Z01-MH-002498). We have no financial disclosures and conflict of interests to disclose.

## Author contributions

S.H. Lee contributed substantially to conception design of work, acquisition, analysis and drafting the manuscript. N.I Cilz contributed substantially conception design of work, acquisition, analysis and interpretation of data for the work, revising critically for important intellectual content. S.K. Williams Avram and A. Cymerblit-Sabba made substantial contribution to the conception and design of the work, revising manuscript critically for important intellectual content. J. Song, K. Courey, A. Howley, M.E. Cooke made substantial contribution to the data acquisition and analysis.

W.S. Young made substantial contribution to design of the work, revising manuscript critically for important intellectual content, final approval of the version to be published.

