## Supplementary for "Stimulation of median raphe terminals in dorsal CA2 reduces social investigation in male mice specifically investigating social stimulus of ovariectomized female mice"

### Supplemental Methods

#### Housing and husbandry

#### Mouse Housing Conditions

All housing and procedures were approved by the Animal Care and Use Committee of the National Institutes of Mental Health. Mice were housed in an AAALAC accredited specific pathogen-free vivarium kept at constant temperature and humidity (~21°C, 50%), in plastic micro-isolator cages (12"x6.5"x5.5") containing wood chip bedding (Nepco Beta-chips) and cotton nestlets. All cages were maintained on high-density ventilated racks (Super Mouse 750, Lab Products Inc.). Mice were maintained on a 12-h light cycle (lights off at 1500h) with *ad libitum* access to standard mouse chow (Purina Lab Diet, Product #5R31) and water bottles. Cages were changed on a bi-weekly basis primarily by the same animal caretaker. All breeding pairs were fed a high-fat diet (Purina Lab Diet, Product #5058) as a measure to reduce pregnancy and pup loss. All offspring were weaned at ~21 days of age into cages with their same-sex littermates. No animals that were singly housed during adolescence were used in behavioral testing.

#### Subjects

Stimulus mice used in social recognition testing were sexually naïve, ovariectomized Balb/c adult females. They were purchased from Jackson Labs (Stock # 000651) and ovariectomized. Briefly, a small dorsal midline incision was made, the muscle wall spread using forceps, and the ovaries were removed. Following a two-week recovery period, females were singly housed for at least one week prior to testing.

For electrophysiology studies, vasopressin-1b receptor Cre knockin (Avpr1b-Cre) mice generated in our lab (1) were used to target CA2 neurons. Cre-dependent TdTomato reporter mice (Ai9) were purchased from the Jackson Laboratory (Strain Name: B6.Cg-Gt(ROSA)26Sortm9(CAG-tdTomato)Hze/J, Stock Number: 007909). These mice were bred to Avpr1b-Cre$^{+/-}$ mice, resulting in Avpr1b-Cre$^{+/-}$;TdTomato double transgenic offspring.

**Genotyping**

**Raphe-cre**

Mice were genotyped by PCR using DNA extracted from tail snips. Raphe-cre mice were genotyped using a forward primer, Cre.m35 (AGAACCTGAAGATGTTCGCGATTATCTTCTATATC) and reverse primer, Cre.c33 (TAGTTACCCCCAGGCTAAGTGCCTTCTCTACAC) with PCR product of 460 bp. PCR was carried out for 40 cycles with denaturation at 95°C, annealing at 65°C, and extension at 72°C, all for 45s.

**Vasopressin-1b receptor Cre knockin (Avpr1b-Cre)**

A single forward primer, V1BR#9 (GAAACGGCTACTCTCTCCGATTCCAAAAGAAAG), was designed for amplification of both WT and recombined loci. The first reverse primer, V1BR#5 (ACCTGTAGATATTTGACAGCCCGG), was designed to amplify the WT loci (762 bp PCR product). The second reverse primer, Cre.c35 (GATATAGAAGATAATCGCGAACATCTTCAGGTTCT), was designed to detect the Cre recombinase transgene (679 bp). PCR was carried out for 40 cycles with denaturation at 94°C, annealing at 60°C, and extension at 72°C, all for 1 min.

**Cre-dependent tdTomato reporter mice (Ai9)**

tdTomato mice were genotyped for the wildtype band using a forward primer, oIMR9020, (AAGGGAGCTGCAGTGGAGTA) and reverse primer, oIMR9021, (CCGAAAATCTGTGGGAAGTC). The mutant band was genotyped using the forward primer, oIMR9105, (CTGTTCCTGTACGGCATGG) and reverse primer, oIMR9103(GGCATTAAAGCAGCGTATCC). PCR was carried out for 40 cycles with denaturation at 95°C, annealing at 65°C, and extension at 72°C, all for 45 s

**Stereotaxic surgery, viral tracing, and optogenetic surgical implantation**

The recombinant adeno-associated virus (AAV) vectors were serotyped with AAV2 coat proteins and packaged by the University of North Carolina Vector Core (Chapel Hill, NC, USA) or Addgene (MA, USA). Male mice (10-12 weeks old) were anesthetized with 100 mg/kg ketamine (100 mg/mL) and 50 mg/kg xylazine (20 mg/mL) and placed on a stereotaxic apparatus. The head was leveled and a small incision was made to expose the skull for localization of the reference lambda and bregma sutures. Burr holes were drilled for subsequent viral injections or optic fiber placements and dental cement was used to hold the optic fiber placements.

**Viral tracing**

For anterograde viral tracing, Raphe-cre mice were injected with Cre-dependent rAAV2-CAG-FLEX-GFP (Addgene, MA, USA) (500 nL of 2 x $10^{12}$ transducing units per ml), serotype 2, in the MR (ML:0.00 mm, AP: -4.2 mm, DV: -4.5 mm) and incubated for 4 weeks.

**Optogenetic Surgical Implantation**

For optogenetic behavioral studies, Cre-dependent AAV-DIO-EF1α-hChR2(H134R)-mCherry (Addgene, MA, USA) (500 nL of 1.5 x $10^{12}$ transducing units per ml), serotype 2, was injected into the MR nucleus (ML: 0.00 mm, AP: -4.2 mm, DV: -4.5 mm). For optical fiber implantation in the dCA2, optical fibers were placed bilaterally at AP: -2.18 mm, DV: -1.67 mm, ML: ±2.56 mm; and secured with dental cement as previously described in Smith *et al*. (2). At the end of behavioral studies, brains were collected, and animals without viral expression and with lose, mis-targeting of fiber optics in stimulated Raphe-cre and or WT mice group were excluded from the analysis.

**Histology and immunohistochemistry**

Brains were processed with immunohistochemistry to detect cells expressing 5-HT and GFP in tissues. Mice were deeply anesthetized and transcardially perfused with 0.9 % saline with 10 U/mL of heparin (heparin lithium salt, Cat. #H-0878, Sigma, St Louis, MO, USA), followed by 4% paraformaldehyde (Cat. # 19210, EMS, Hatfield, PA, USA) in phosphate buffer (PB), pH 7.4, and incubated for 24 hours. Brains were stored in 20% sucrose in PB at 4°C for at least 48 hours. Brains were then mounted with O.C.T compound (Tissue-Tek, Torrance, CA, USA) at -20°C, and cut into 40 μm coronal sections using a cryostat and stored in cryoproctectant (30% sucrose in PB, pH 7.4, and 30% ethylene glycol) at -20°C. Sections were washed in PB, pH 7.4, and used for immunohistochemistry.

Transgenic Raphe-cre x B6.129(Cg)-Gt(ROSA)$26^{Sortm4Luo/J}$ GFP brain sections were used for free-floating immunohistochemistry to detect cells expressing GFP. Sections were incubated in blocking solution (PB, pH 7.4, with 0.2 % Triton X100 and 10 % normal goat serum) for 30 min at room temperature (RT). Sections were then incubated with an anti-GFP antibody ( #AB8370, Takara Bio USA, Mountain view, CA, USA) in 1:500 dilution overnight at 4°C. Sections were washed in 24-well plate in PBST (phosphate-buffered saline, pH 7.4, with 0.01% triton-X100) solution on a 100 RPM orbit shaker for 10 min, 3 times at RT. Sections were incubated with a secondary antibody, Cy-2 affinity pure goat anti-rabbit antibody (Jackson ImmunoResearch Laboratories Inc., West Grove, PA, USA), in 1:250 dilution for 1 hour at RT. Sections were washed in 24-well plate in PBST, pH 7.4, solution on a 100 RPM orbit shaker for 10 min, 3 times at room temperature.

Brain sections from Raphe-cre mice injected with Cre-dependent rAAV2-CAG-FLEX-GFP were stained for 5-HT expression using free-floating immunohistochemistry. Sections were incubated in blocking solution (PB, pH 7.4, with 0.2 % Triton X100 and 10 % normal goat serum) for 30 min at RT. Sections were then incubated with anti-5-HT rabbit antibody (cat #20080, ImmunoStar, Hudson, WI, USA) in 1:1000 dilution at 4°C overnight. Sections were washed in 24-well plate in PBST, pH 7.4, solution on a 100 RPM orbit shaker for 10 min, 3 times at RT. Sections were incubated with secondary goat anti-rabbit Alexa Fluor 647 antibody in 1:250 dilution at RT for one hour. Slides were stained with 300 nM of 4'6-diamidino-2-phenylindole (DAPI) for 2 minutes and washed with 1xPBS.

Images were acquired using a fluorescent microscope (Nikon 50i) with a DAPI filter for DAPI-stained sections, a fluorescein isothiocyanate (FITC) filter for GFP-expression sections, and a tetramethylrhodamine isothiocyanate (TRITC) filter for mCherry fluorescence. Images were processed using iVision (BioVision Technologies) and ImageJ (U.S. National Institutes of Health).

**Electrophysiology**

Whole-cell recordings were obtained from coronal slices (350 μM) prepared from Raphe-Cre mice (9 weeks old) after 5-12 week viral infection. To prepare slices, the animal was deeply anesthetized with isoflurane and the brain was rapidly removed into a cutting solution containing (mM): 248 mM sucrose, 10 mM glucose, 10 mM MgCl2, 1 mM CaCl2, 26 mM NaHCO3, 1 mM KCl. After slicing, the sections recovered for 20-30 minutes 33 C in a recovery solution containing (mM): 90 mM sucrose, 80 mM NaCl, 10 mM glucose, 25 mM NaHCO3, 3.5 mM KCl, and 1.25 mM NaH2PO4. After recovery, slices were transferred to a holding chamber containing the following solution (mM): 97 NaCl, 2.5 KCl, 1.25 $NaH_2PO_4$, 30 $NaHCO_3$, 20 HEPES, 3 Na-Pyruvate, 5 Lasorbic acid, 2 $CaCl_2$, 2 $MgCl_2$, and 25 glucose that was continuously oxygenated (95%$O_2$/5%$CO_2$). For recordings, slices were transferred to a recording chamber mounted on a Nikon FN1 upright microscope equipped with infrared differential interference contrast (DIC) and epifluorescence imaging optics and perfused (~1-2 mL/min) with an artificial cerebral spinal fluid solution containing (mM): 130 NaCl, 24 $NaHCO_3$, 3.5 KCl, 1.25 $NaH_2PO_4$, 2 $MgCl_2$, 2 $CaCl_2$, and 10 glucose continuously oxygenated (95%$O_2$/5%$CO_2$). Current-clamp recordings were acquired using pClamp 11 software (sampling rate = 10 kHz) interfacing with a Multiclamp 700B amplifier (Molecular Devices). Patch electrodes (3-6 MΩ) were filled with (mM): 100 K-gluconate salt, 0.6 EGTA, 5 MgCl2, 8 NaCl, 2 ATPNa2, 0.3 GTPNa, 7 Phosphocreatine, and 33 HEPES. Light-induced responses (1 ms pulse) were evoked via a collimated LED (Cool LED) mounted to the microscope and triggered with pClamp. For light-evoked EPSCs onto interneurons, 20 sweeps were collected both with and without LED stimulation. Peak negative-going currents were analyzed in Clampfit in both conditions. For spontaneous inhibitory postsynaptic current (sIPSC) recordings, we used slices prepared from offspring of Avpr1b-Cre mice crossed with an Ai9 reporter line. We patched tdTomato-positive neurons of the CA2 region using an internal solution containing (mM) 70 gluconic acid, 35 CsCl, 0.3 EGTA, 1 MgCl2, 4 ATPNa2, 0.3 GTPNa, 10 phosphocreatine, and 10 HEPES (pH adjusted to 7.4 with CsOH). The membrane potential was clamped at -65 mV and the external solution was supplemented with ionotropic glutamatergic blockers (10 µM DNQX and 50 µM APV) to isolate GABAergic events. The sIPSC events were detected in Clampfit using an event detection template search and data was further analyzed in Origin 2018. For interneuron recordings, cells were patched in the stratum radiatum and stratum lacunosum moleculare. The average resting membrane potential of the last minute of 5-HT application was compared to the average resting membrane potential of the baseline period.

**Behavioral Analysis**

Videos of behavioral tests were recorded from above and coded by an observer blind to the identity of the mouse using Noldus Observer v12. The total duration and frequency of behavior is reported. The behavioral tests were conducted in a week apart from each test.

**Social recognition (SRM) test**

Duration of sniffing was manually scored, and experimenter was blinded to the subject. Sniffing was only recorded when the experimental mouse initiated the behavior. When that mouse contacted the stimulus mouse with its nose, a sniffing bout was initiated and when nose of experimental mouse was no longer in contact, the bout was ended.

**Object recognition**

Duration of sniffing was manually scored, and experimenter was blinded to the subject. When experimental mouse contacted the object with their nose, a sniffing bout was initiated and when nose of experimental mouse was no longer in contact, the bout was ended.

**Social interaction test**

Duration of sniffing was manually scored, and experimenter was blinded to the subject. When experimental mouse contacted the stimulus mouse or object with their nose, a sniffing bout was initiated and when nose of experimental mouse was no longer in contact, the bout was ended. Chamber and objects were cleaned thoroughly with 70% ethanol between trials.

**Marble Burying Test**

Latency to begin digging around the first marble was manually scored, and experimenter was blinded to the subject.

**Open-field test**

For locomotor activity, distance travelled was measured. Center was defined on Ethovision as the inner 50% zone of the open-field arena, center duration as the time spent in the inner zone, and center frequency as the frequency to enter inner zone.

**Supplemental Figures**

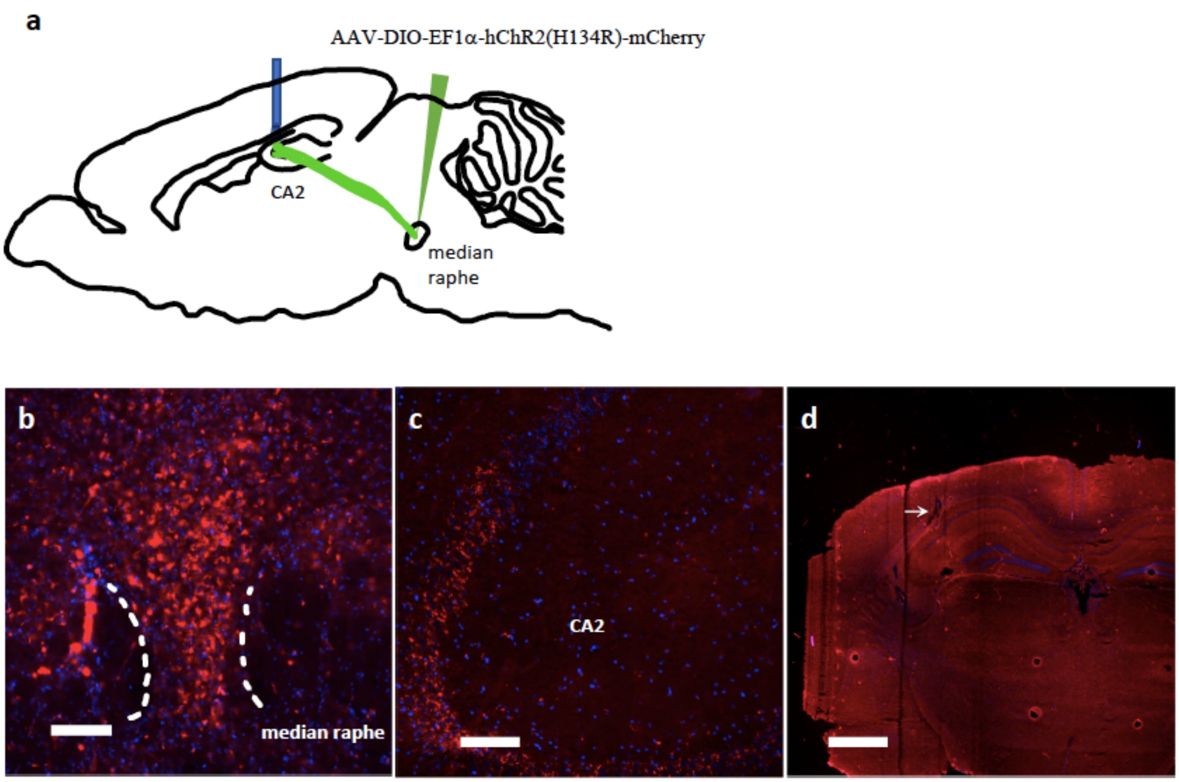


**Supplemental Figure 1.** (a) Transgenic Slc6a4 Raphe-cre mice were injected with AAV-EF1a-DIO-hChR2(H134R)-mCherry in the MR region. mCherry expression is in (b) MR neurons and (c) fibers projecting to the dCA2. (c) The optic fiber probe was placed just above the dCA2 region. Scale bars: (b, c) 100 μm, (d) 1000 μm.

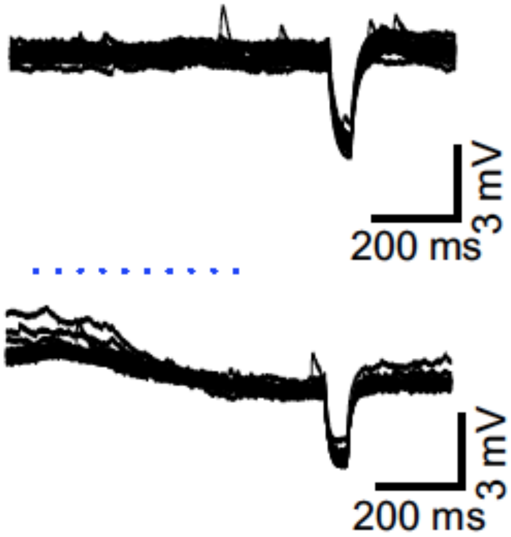


**Supplemental Figure 2.** We also observed 1 cell in the SR, proximal to CA2, which showed a membrane hyperpolarization in response 10 light pulses delivered at 20 Hz.

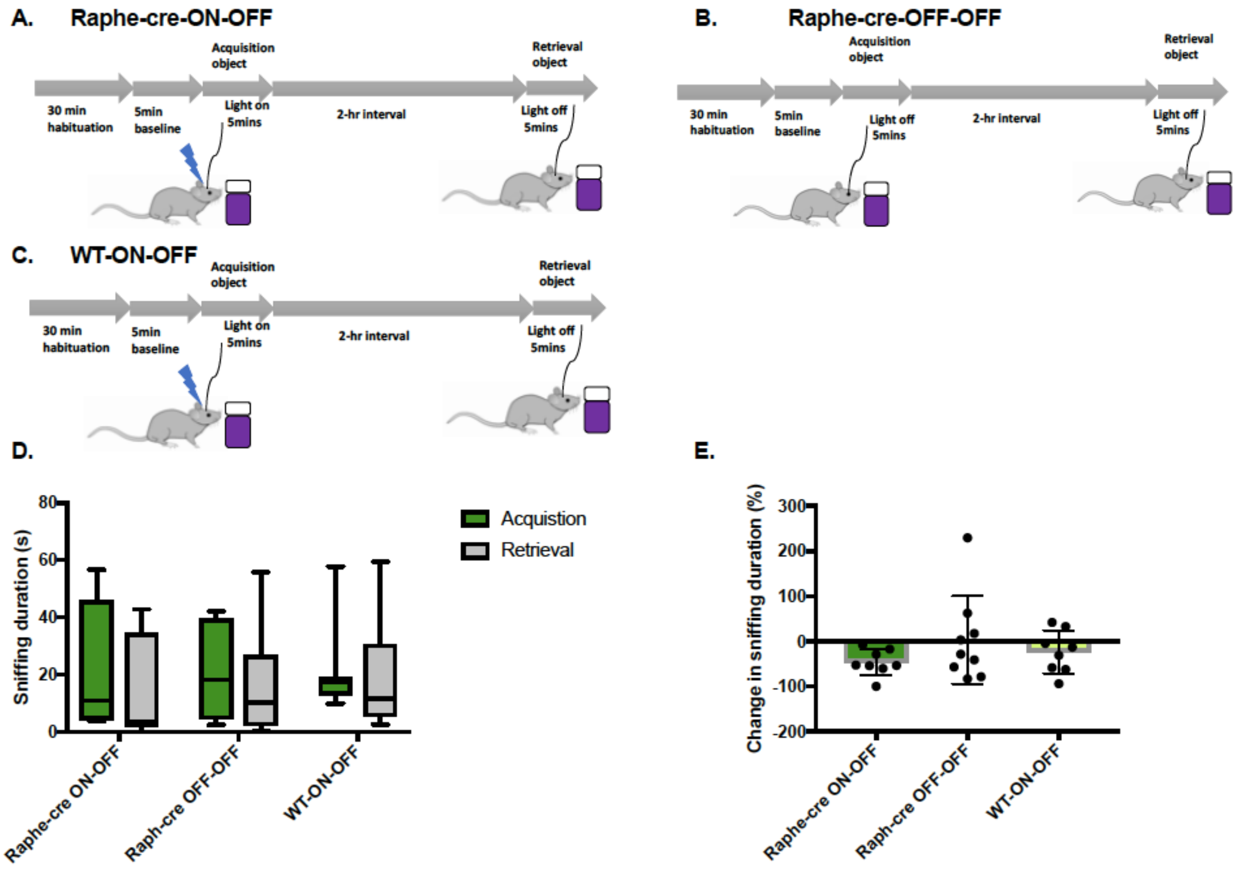


**Supplemental Figure 3. Object recognition memory test with 2-hour interval.** (A) Illustrates Raphe-cre mice optogenetically stimulated during acquisition period (n = 8). (B) Non-stimulated Raphe-cre group (n = 9). (C) WT mice optogenetically stimulated during acquisition period (n = 8). (D) Object sniffing duration during acquisition and retrieval period. (E) Change in sniffing duration of an object calculated by $(\text{Retrieval sniffing duration} - \text{Acquisition sniffing duration}\}/\text{Acquisition sniffing duation} \times 100$. There were no significant differences between the genotypes and stimulations.
